# Insight into the gut virome in patients with multiple sclerosis

**DOI:** 10.1101/2023.11.17.567435

**Authors:** Suresh C Bokoliya, Jordan Russell, Hanshu Yuan, Zongqi Xia, Laura Piccio, Yanjiao Zhou

**Affiliations:** Department of Medicine, University of Connecticut Health Center, Farmington, CT 06030, USA; Department of Neurology, University of Pittsburgh, Pittsburgh, PA, USA; Charles Perkins Centre and School of Medical Sciences, The University of Sydney, Camperdown, NSW, Australia

**Keywords:** RRMS, MDA, VLP, Virome

## Abstract

Multiple sclerosis (MS) is an autoimmune condition associated with dysbiosis in the bacterial element of microbiome, yet limited information exists regarding dysbiosis in the virome. In this study, we examined the virome in 20 relapsing-remitting MS (RRMS) patients and 22 healthy controls (HC). We extracted virus-like particles (VLP) genomic DNA through sequential filtration, followed by deep metagenomic sequencing approaches with and without multiple displacement amplification (MDA). We found significantly lower diversity in the gut virome of RRMS patients relative to HC, consistent across both sequencing methods. MDA method identified reduced relative abundance of *Microviridae* and *Myoviridae* bacteriophage, and eukaryotic virus such as *Herpesviridae* and *Phycodnaviridae* in RRMS patients compared to HC. Non-MDA methods showed reduction in relative abundance of *Siphoviridae* bacteriophage and eukaryotic viruses such as *Ackermannviridae*, *Demerecviridae*, *Dicistroviridae*, *Herelleviridae*, *Mesnidovirineae* in RRMS patients. Cluster analysis revealed that the whole virome family was dominated by *Podoviridae* and *Siphoviridae* clusters. Comparing dietary metadata between these clusters, RRMS patients in the *Siphoviridae*-dominated Cluster B showed significantly higher consumption of refined grains and salad dressings compared to those in the *Podoviridae*-dominated Cluster A. Correlation analysis between gut viruses and bacteria demonstrated that *Siphoviridae* exhibited positive correlations with many different bacterial genera. Conversely, *Microviridae* displayed negative correlations with many different bacterial genera. These findings underscore the alterations in viral diversity and taxonomic composition of the gut virome in RRMS patients. Our study represents the first step in understanding the gut virome in MS patients, providing a groundwork for future research on the role of the gut virome in the context of MS.

## Introduction

Multiple sclerosis (MS) is a complex autoimmune disease of the central nervous system characterized by demyelination. Globally, MS affects approximately 2 million individuals, and currently, there is no cure available. The most prevalent form of MS is relapsing-remitting MS (RRMS) that affects 85% of the total patient population (1). While the exact cause is unknown, a combination of genetic and environmental factors is believed to contribute to its development (2). Several studies have shown that changes in the gut microbiome can influence the immune system, affect the stability and function of the blood-brain barrier, initiate demyelination, and interact with various immune cell types in MS (3). As a novel and essential player in immune and metabolic homeostasis, gut microbiome has become a potential therapeutic target in MS (4). However, most studies on the gut microbiome in MS have primarily focused on bacteria, neglecting the reservoir of diverse viruses that also reside in the gut known as the “gut virome”. The gut virome is a dynamic entity that can interact with the gut bacterial microbiota, impacting the host’s immune response and overall health. In a healthy gut, the gut virome remains stable over time but diversifies during illness (5). This diversification is also modulated by individual dietary patterns (6, 7), ethnic disparities, geographic factors, and urbanization trends (8). The gut virome predominantly comprises bacteriophages, numbering approximately 1×10^8^ to 2×10^9^ (9), a count parallels that of gut bacteria (10). Bacteriophages play a pivotal role in gut physiology by exerting influence on the intestinal lining by adhering to mucus (11). In addition, a diverse range of eukaryotic virtues are also populated in the gut (12).

While studies have shown dysbiosis of gut virome in multiple diseased conditions including inflammatory bowel diseases (13–15), colon cancer(16), diabetes (17–20), alcoholic and non-alcoholic liver ailments (21, 22). Notably, few studies conducted among patients with autoimmune disease such as rheumatoid arthritis (23, 24) and systemic lupus erythematosus (24, 25), with only a recent study shedding light on alterations in the gut virome related to MS (26). Specific viruses, such as Cytomegalovirus, Varicella zoster virus and Human Endogenous Retrovirus-W and Epstein-Barr virus, have been implicated in development of MS and immune modulation (27). Whether the gut virome is associated with MS has not been explored. Technical challenges such as the absence of an universal taxonomic marker for virome sequencing and the requirement for viral enrichment techniques (28) have impeded our understanding of the role of the virome in diseases. Viral metagenomic sequencing, while more complex, offers a more comprehensive representation of viral communities and enhanced detection sensitivity for rare and low-abundant viruses (29).

To gain insight into the gut virome profile of RRMS patients and their healthy controls, we performed viral enrichment using sequential filtration to isolate virus-like particles (VLP), followed by deep metagenomic shotgun sequencing with and without multiple displacement amplification (MDA). We identified a complex DNA viral community comprising of bacteriophage and eukaryotic viruses and showed alteration of the gut virome in RRMS patients compared to health controls. We have identified multiple clinical and dietary factors associated with the gut virome. Our study provides the first insight into the gut virome in MS patients, laying the foundation to study the role of the gut virome MS in future.

## Methods

### Subjects

We conducted an analysis of the clinical characteristics of 42 subjects, consisting of 20 individuals diagnosed with RRMS and 22 healthy controls (HC). All participants were enrolled at the Washington University School of Medicine in St. Louis City, Missouri, U.S.A. The individuals with RRMS were diagnosed by clinicians based on the McDonald 2010 criteria (1). Prior to their involvement, all subjects provided written informed consent. The demographic and clinical characteristics of these subjects are presented in **Table 1**.

**Table 1:** Demographics and clinical characteristics of RRMS and HC.

| **Group** | **RRMS** | **HC** | ***p*-value** |
| --- | --- | --- | --- |
| Number of subjects | 20 | 22 | *p*>0.05 |
| Age in years at sample collection [average] | 30.5-56.3 [38.8] | 30.5-56.5 [38.4] | *p*>0.05 |
| BMI [average] | 17.8-49.8 [28.1] | 17.6-40.5 [27.0] |  |
| Beck depression score [average] | 0-30 [7.6] | 0-17 [3.6] | *p*=0.054 |
| Gender<br>Male:Female:NA | 3:18:1 | 2:18:0 | *p*>0.05 |
| Race | White-18<br>Black/African-American-2 | White-16<br>Black/African-American-4<br>NA-2 | *p*>0.05 |
| Antibiotic use<br>Yes:No:NA | 1:17:2 | 1:20:1 | *p*>0.05 |
| Use of steroids<br>Yes:No:NA | 2:14:2 | 0:17:5 | *p*>0.05 |
| Use of probiotic<br>Yes:No:NA | 2:15:3 | 2:16:4 | *p*>0.05 |
| Use of supplements<br>Yes:No:NA | 10:7:3 | 6:14:2 | *p*>0.05 |
| Constipation<br>Yes:No:NA | 4:13:3 | 2:18:2 | *p*>0.05 |
| Diarrhea Yes:No:NA | 2:14:4 | 1:19:2 | *p*>0.05 |
| Age at MS diagnosis [average] | 18.4 – 52.7 [33.9] |  |  |
| Years after at MS diagnosis [average] | 0 – 23.1 [4.6] |  |  |

RRMS: Relapsing-Remitting Multiple Sclerosis; HC: Healthy controls

### Stool collection

Stool samples were self-collected and carefully placed on frozen gel packs, then shipped overnight to the research laboratory. Upon arrival at the laboratory, the stool samples were promptly stored at −80°C until further procedures were carried out. To ensure consistency and minimize any potential batch effects, stool samples collected at the baseline and six-month time points were processed simultaneously for DNA extraction and microbiome sequencing.

### Bacterial DNA isolation and sequencing

We employed 16S rRNA gene sequencing protocol as described in our previous report (30). Stool DNA extraction was carried out using the Qiagen Power Soil DNA Extraction kit (Qiagen, MD, U.S.A). For 16S rRNA gene sequencing, the hyper-variable regions V1–V3 of the 16S gene were amplified using primers 27F and 534R (27F: 5′-AGAGTTTGATCCTGGCTCAG-3′ and 534R: 5′-ATTACCGCGGCTGCTGG-3′). Subsequently, amplified libraries were prepared and sequenced on the Illumina MiSeq platform using a V3 2 × 300 bp paired end sequencing protocol, targeting a read depth of 10,000 reads per sample. Initial processing of raw sequencing data was managed by Illumina’s software (Illumina, CA, U.S.A). Sample deconvolution was conducted with one mismatch allowed in primers and zero mismatches in barcodes. Further data processing involved the removal of sequences with low quality (average quality score <35) and sequences with ambiguous bases. Chimeric amplicons were identified and removed using UChime (v4·2·40), and amplicon sequences were clustered into operational taxonomic units (OTUs) using Usearch against the SILVA_132_SSURef_Nr99 database at a 97% threshold. Taxonomic assignment was performed using RDP-classifier (v2·11) with a 0.8 confidence value cutoff. To ensure data integrity, potential contaminant taxa found in DNA extraction controls and PCR controls were exluded from the sequenced samples. Sequencing and data processing were conducted at The Jackson Laboratory for Genomic Medicine, CT, U.S.A.

### VLP Preparation, Viral genomic DNA isolation, and sequencing

VLPs were isolated by initially mixing approximately 1.0 gram of stool with 10 mL of SM Buffer which was adjusted to a pH of 7.5 (Thermo Fisher Scientific, MA, U.S.A). Subsequently, the samples were vigorously vortexed for 5 minutes at 3000 RPM using Vortex Genie 2 (Scientific Industries, INC, NY, U.S.A). After this step, the samples were subjected to centrifugation at 4000xg for 10 minutes at 4°C. The resulting supernatants were filtered through a 0.45 µM PVDF filter SteriFlip Unit (Milipore Sigma, MO, U.S.A) followed by a 0.22 µM polyethersulfone filter SteriFlip Unit (Milipore Sigma, MO, U.S.A). The filtered VLP supernatants were adjusted to a 10 mL volume using sterile SM Buffer. The VLPs were subsequently concentrated down to 180 µL using Amicon Ultra-15 100K Centrifugal Filter Units (Milipore Sigma, MO, U.S.A). To remove any free contaminating nucleic acids, the concentrated VLPs were treated with 4 µL of Ambion Turbo DNAse (equivalent to 8 Units) and 0.5 µL of Ambion RNase 1 (equivalent to 50 Units), both sourced from Invitrogen, Thermo Fisher, MA, U.S.A. Additionally, 20 µL of a buffer containing 10 mM CaCl2 and 50 mM MgCL2, obtained from Millipore Sigma, MO, U.S.A was incorporated in this treatment process. DNase/RNase treatment was deactivated by incubating the samples at 70°C for 10 minutes.

DNase/RNase treated VLPs were used for genomic DNA extraction using Qiagen QIAamp DSP Virus Spin Kit (Qiagen, MD, U.S.A) following the manufacturer’s protocol, with specific modifications such as omitting of carrier RNA, resuspending Qiagen Protease Buffer in Buffer, and the eluting resulted DNA in 40 µL of DNase/RNase Free water (Thermo Fisher Scientific, MA, U.S.A).

We employed MDA method to generate viral libraries for metagenomic whole-genome shotgun sequencing (mWGS). In the MDA method, VLP genomic DNA underwent amplification using the Illustra GenomiPhi V2 DNA Amplification Kit (formerly GE Healthcare Life Sciences but now Cytiva, Tokyo, Japan) following the manufacturer’s instructions. The MDA-amplified VLP DNA was then processed using the Zymo Genomic DNA Clean and Concentrator-10 kit (Zymo Research, CA, U.S.A) and eluted in Zymo DNA Elution Buffer. Library preparation for VLP genomic DNA involved using >25ng input DNA with the Nextera DNA Flex Library Preparation Kit (currently named Illumina DNA prep, Illumina, CA, USA). Deviating from the Illumina protocol’s full reaction volume (50 µL), we opted for a quarter reaction volume (12.5 µL), reducing all components, including DNA concentration and volumes, by one-fourth. Dual-indexed paired-end libraries with an average insert size of 350bp were created from VLP genomic DNA. The bead-linked transposomes in the Nextera DNA Flex kit facilitated DNA fragmentation and the tagging of Illumina sequencing primers. Subsequently, double-sided cleaned tagged libraries were sequenced on NovaSeq 6000 (Illumina, CA, USA) using a 2 × 250 bp paired end sequencing run, targeting 50M reads/sample. For comparison, we also include a routine library preparation without MDA. Approximately <25ng VLP genomic DNA served as input for library preparation.

The raw sequencing reads were processed as we have done previously (30). In brief, we demultiplexed the raw reads, and further processed them by (a) removing human reads using NCBI’s BMTagger (v3·101) (ftp://ftp.ncbi.nlm.nih.gov/pub/agarwala/bmtagger); (b) removing duplicated reads using GATK-Picard 4·1·0 (MarkDuplicates); (c) trimming low-quality bases and low-complexity screening using PRINSEQ (v0·20·4). We further removed bacterial reads by aligning the reads to NCBI bacterial database. The resulting reads were aligned to NCBI virus database using BWA software. The final reads were assigned to different taxonomy based on NCBI viral taxonomic classification using EFetch (30) and TaxonKit (31). The viral hits in different samples were summarized into a table. To account for sequencing depth variations across samples, viral hit counts were normalized using the trimmed mean of M-values (TMM) method implemented in the edgeR package in R (32), separately for MDA and non-MDA counts. Contaminants were removed by excluding viral hits present in any of the negative controls, with the exception of crass-like phage. The resulting normalized and filtered counts were converted into relative abundance, and low relative abundance reads were filtered to retain hits with an average relative abundance exceeding 0.01%.

### Diet record

Subjects recorded a four-day food diary covering two weekends and two weekdays, provided insights into their diets (30, 31). The Nutrition Coordinating Center (NCC) Food Group Serving Count System was used to analyze the diaries, estimating food group consumption (32). The study focused on eighteen food groups, including daily servings of yogurt with live active cultures, whole grains, vegetable, vegetable oils, sugar sweetened soft drinks, salad dressings, refined grains, poultry, plain and flavored cow’s milk, nut and seeds, meat, fruits, fish and shellfish, eggs, dairy cheese, cold cuts and sausages, butter and animal fats, as well as beer, liquor, and wine. The daily serving sizes for each food group were determined based on the 2000 Dietary Guidelines for Americans (33). For foods not covered by these recommendations, serving sizes as defined by the United States Food and Drug Administration (FDA) were used.

### Data analysis

In our analysis, we individually examined demographic and clinical variables between RRMS and HC within the PERMANOVA model. It’s important to note that the variance explained by each variable was calculated independently, without considering the influence of other variables. For virome composition analysis, the viral count tables were combined with associated metadata using phyloseq (34). Categorical metadata variables were compared between diagnosis groups using Fisher’s exact test, while continuous metadata variables were assessed using the Wilcoxon test. Alpha diversity metrics, including richness and Shannon diversity, as well as virus prevalence, were computed using the Microbiome package in R. Overall differences in virome (beta diversity) among enrichment methods and diagnosis groups were evaluated using PERMANOVA through the adonis function in the vegan package and visualized via principal coordinate analysis (PCoA). Differential relative abundance analysis between diagnosis groups was performed using Linear discriminant analysis Effect Size (LefSe) (35). Spearman correlations analysis was performed to investigate potential associations between dietary factors and phage relative abundance. Various data visualizations, such as barplots, boxplots, scatterplots, and heatmaps, were generated using ggplot2 (36). Metagenome-assembled genomes (MAGs) of viruses were constructed using metaSPAdes v3.14.18 (37). A network was constructed using Cytoscape (38) and analyzed to explore the interactions between bacterial and viral communities dominated RRMS and HC. Spearman correlation was used to identify significant correlation between virus-bacteria network.

## Results

### Demographics and clinical characteristics of RRMS and healthy controls

There were no significant differences observed between participants with RRMS (n=20) and HC (n=22) in categorical variables such as race, sex, reports of constipation or diarrhea, or the use of antibiotics, steroids, probiotics, and/or supplements at the time of sampling (Fisher’s exact test, *p*>0.05). Similarly, no significant differences were found in continuous variables, including age at sample collection, body-mass index (BMI), and Beck depression score (Wilcoxon test, *p*>0.05). Although the Beck depression score was slightly higher among RRMS subjects (average: 7.61) compared to HC (average: 3.62), this difference was marginally insignificant (*p*=0.054). Overall, there were no significant clinical variables that differed between HC and RRMS (**Table 1**).

### Virome detection in stools by MDA and non-MDA methods

Samples from MDA method yielded significantly fewer reads than non-MDA (Wilcoxon, *p*=3.9E-14). However, MDA samples exhibited a notably higher proportion of viral reads, reaching an average of 68.9% with a median value of 99.7%. In contrast, non-MDA samples contained a lower proportion of viral reads, accounting for only 31.8% of the assigned reads on average, with a median of 19.4%. In addition, after filtering for contamination identified from negative controls and low-relative abundance sequences (<0.01%), we found 140 viral hits unique to samples with MDA and 262 viral hits unique to samples with non-MDA method (**Fig. 1A**). Additionally, 110 viral hits were shared between MDA and non-MDA samples. These findings indicates that different method strongly influence viral detection, and the combination of these methods provides a comprehensive view of the gut virome.

**Fig. 1.**
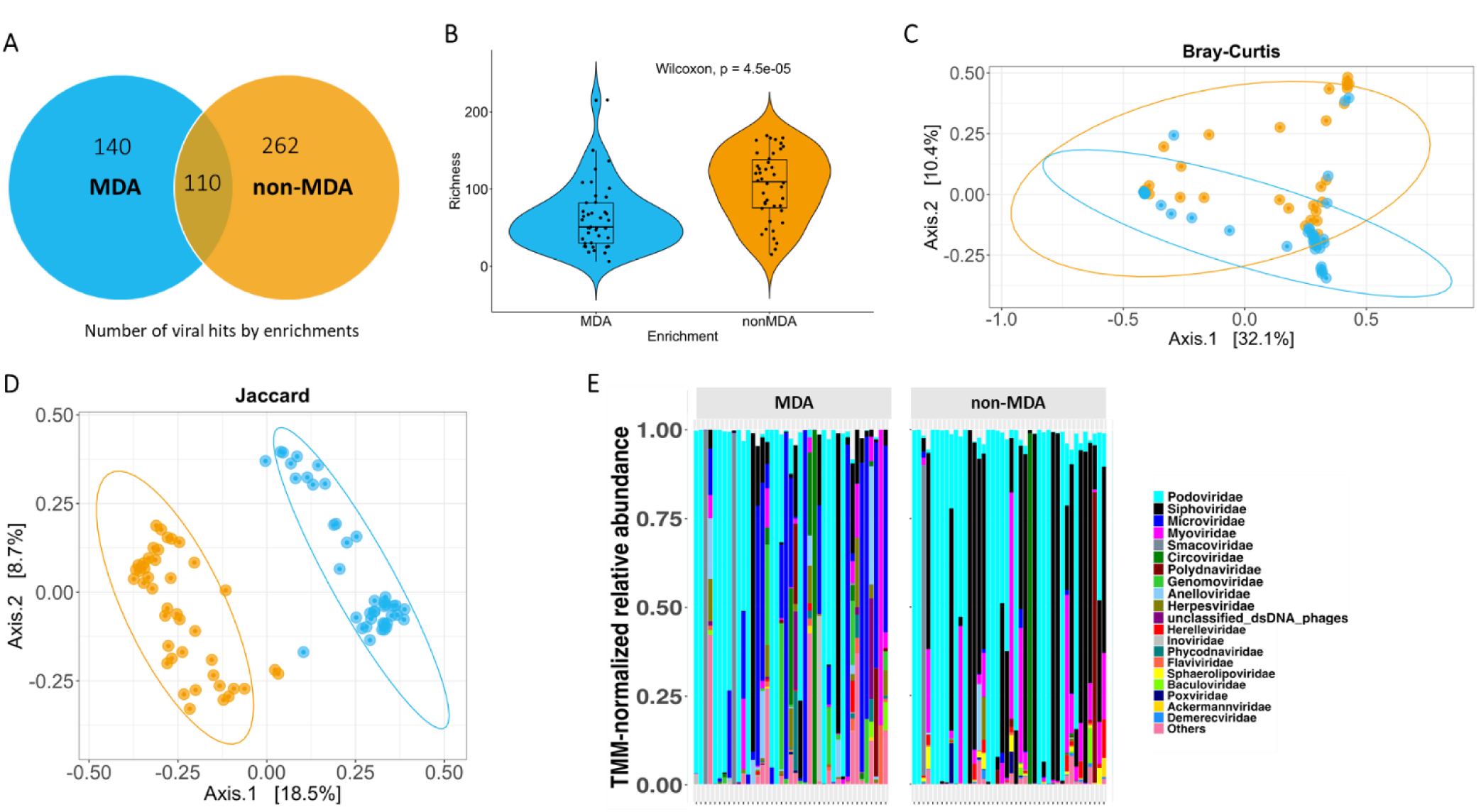
illustrates the comparison of gut virome characteristics and relative abundance between MDA and non-MDA methods. **A**: The Venn diagram provides a comparison of viral hits, indicating both shared and unique ones, observed through MDA and non-MDA methods; **B**: Violin and box plots depict virome richness using MDA and non-MDA methods. Data obtained from the MDA approach is represented in sky blue, while data from non-MDA is shown in orange. The violin plot showcases the kernel probability density of the data across various values, while the box plot presents the data in quartiles, with the median indicated by a horizontal line within the box and the “box” representing the middle 50% of the data. The upper and lower whiskers denote the maximum and minimum values, respectively. The viral richness is significantly different using a Wilcoxon test, and the *p*-values are provided; **C**: Bray-Curtis and **D**: Jaccard distances are presented to compare viral communities between MDA and non-MDA methods. The sky-blue ellipse represents MDA method, while the orange ellipse represents non-MDA method. Colored ellipses denote 95% confidence intervals, and the percentage of variability explained by each axis is displayed; **E**: A stacked bar plot displays the top 20 most abundant gut viruses, distinguishing between MDA and non-MDA approaches.

MDA method resulted in a significant decrease (Wilcoxon, *p*=4.5E-05) in viral richness compared to non-MDA (**Fig. 1B**). However, Shannon viral diversity did not differ significantly between MDA and non-MDA samples (Wilcoxon test, *p*=0.19). This suggests that the virome’s overall evenness remained stable in both MDA and non-MDA samples. Both methods also significantly affected the overall viral community structure, both in terms of viral relative abundance (PERMANOVA, Bray-Curtis, *p*=0.005) and viral presence or absence (PERMANOVA, Jaccard, *p*=0.001). PCoA revealed distinct separation between virome communities based on the MDA and non-MDA (**Fig. 1C** and **1D**).

MDA samples showed higher proportions of circular, single-stranded DNA viruses, predominantly of tail-less bacteriophages from the family of *Microviridae* and increased proportions of eukaryotic viruses such as *Genomoviridae*, *Circoviridae*, and *Anelloviridae* (Fig. 1E). In contrast, non-MDA samples were dominated by tailed bacteriophages with double-stranded DNA genomes, from three families within the *Caudovirales* order, namely *Podoviridae, Myoviridae* and *Siphoviridae* (**Fig. 1E**). Based on these significantly distinct virome profiles, we analyzed the MDA and non-MDA-derived virome separately to compare virome differences between RRMS and HC.

### Virus diversity and composition differ by MS diagnosis

When comparing the virome diversity from RRMS patients with HC, it was observed that the viral Shannon diversity exhibited a significant reduction in RRMS patients (MDA: *p*=0.03 and non-MDA: *p*=0.03). (**Fig. 2A**) MDA and non-MDA methods exhibited no significant change in viral richness between RRMS and HC (MDA: *p*=0.73 and non-MDA: *p*=0.20).

**Fig. 2.**
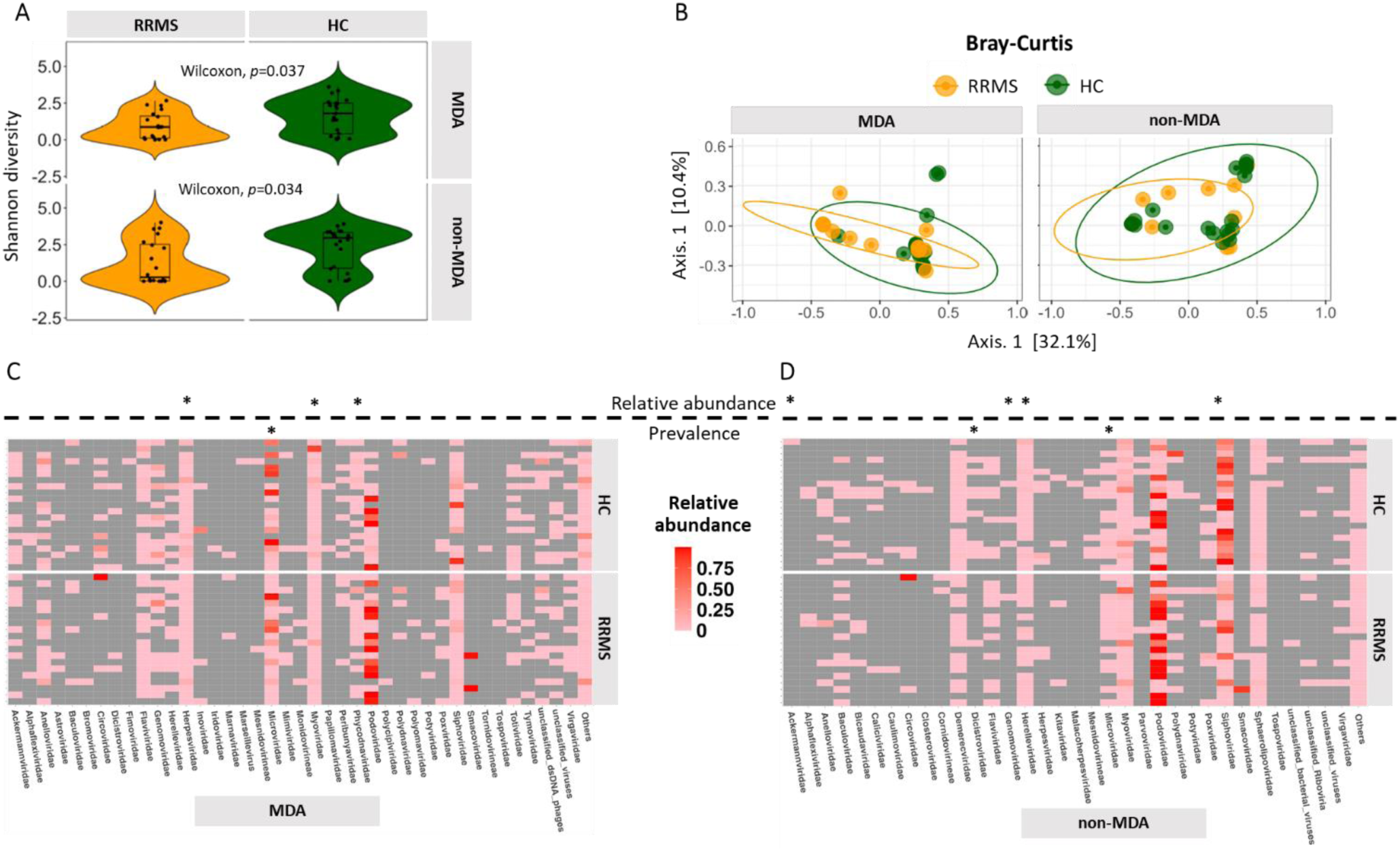
illustrates a comparative analysis of gut virome diversity and composition in individuals with RRMS and HC using both MDA and non-MDA method. **A**: Violin and box plots depict virome diversity when using MDA and non-MDA in both RRMS and HC. Data obtained from the RRMS is represented in orange, while data from HC is shown in green. The violin plot showcases the kernel probability density of the data across various values, while the box plot presents the data in quartiles, with the median indicated by a horizontal line within the box and the “box” representing the middle 50% of the data. The upper and lower whiskers denote the maximum and minimum values, respectively. The virome diversity were statistically compared using a Wilcoxon test, and the *p*-values are provided; **B**: Bray-Curtis distances are displayed to compare viral communities between RRMS and HC using MDA and non-MDA method. The orange ellipse represents RRMS, while the green ellipse represents HC. Colored ellipses denote 95% confidence intervals, and the percentage of variability explained by each axis is displayed; **C** and **D**: A heatmap visually represents the relative abundance of the 35 most prevalent viruses within the virome for both RRMS and HC groups, using both MDA and non-MDA methods. The star symbol (*) is used to denote a significant difference in relative abundance or prevalence of viral strain between individuals with RRMS and HC.

A significant difference (PERMANOVA, Bray-Curtis, *p*=0.037) in the overall virome community structure between RRMS and HC was evident when MDA was employed. In contrast, such a differentiation was not observed when using the non-MDA method (PERMANOVA, Bray-Curtis, *p*=0.061) as depicted in **Fig. 2B**. However, RRMS and HC virome did not differ based on virus presence/absence for either method (PERMANOVA, Jaccard, MDA: *p*=0.854 and non-MDA: *p*=0.591).

Subsequently, we examined the relative abundance of specific viral families that exhibited significant differences between RRMS and HC using both MDA and non-MDA methods. MDA led to the identification of four viral families—*Herpesviridae*, *Microviridae*, *Myoviridae* (the most abundant single-stranded DNA virus), and *Phycodnaviridae* that were significantly reduced in RRMS patients (*p*<0.05 and FDR <0.10) compared to HC (**Fig. 2C**). Using non-MDA enrichment, we identified additional six viral families consist of *Ackermannviridae*, *Demerecviridae*, *Dicistroviridae*, *Herelleviridae*, *Mesnidovirineae*, and *Siphoviridae* that were significantly reduced in RRMS patients (*p* < 0.05 and FDR < 0.10) compared to HC (**Fig. 2D**). Further, the relative abundance of *Myoviridae* also exhibited a decreasing trend in RRMS patients using non-MDA method (*p*=0.08 without FDR adjustment). It’s noteworthy that the *Siphoviridae* family comprises double stranded DNA phages infecting bacteria and archaea as their natural hosts. Due to its high relative abundance in non-MDA samples, we further investigated genus-level differences within the *Siphoviridae* family and identified a significant depletion of *Lactococcus* phages from the *Skunavirus* genus in RRMS patients. Together, our data suggests that RRMS patients exhibit a significant reduction in specific single-stranded and double-stranded DNA viruses, and both MDA and non-MDA methods enhance the sensitivity to detect differentially abundant viral families between RRMS and HC.

### Association of diet, demographics, and clinical factors with virus relative abundance

Dietary choices can have a profound impact on the gut microbiome, including the virome (6). In our study, we found significant correlation between various viral families and key dietary categories, including meat, oil, poultry, fruits, vegetables, and grains using both MDA and non-MDA methods. Notably, we observed a strong negative correlation between the relative abundance of *Baculoviridae* and the consumption of eggs. Additionally, we found that the relative abundance of *Bicaudaviridae* was negatively correlated to whole grain intake and positively correlated with vegetable oil consumption. Furthermore, *Closteroviridae* relative abundance showed a negative correlation with refined grain servings. *Demerecviridae* relative abundance was negatively correlated to fish and shellfish consumption but positively correlated with yogurt containing live and active cultures. Moreover, our study revealed a strong negative correlation between the relative abundance of *Microviridae* and the consumption of eggs, butter, and animal fats. Conversely, a positive correlation was observed between *Microviridae* relative abundance and the consumption of fruits and vegetables. We also noticed a strong negative connection between the prevalence of *Podoviridae* and the consumption of salad dressings, while the consumption of salad dressing was positively correlated with the relative abundance of *Siphoviridae*. Additionally, *Polydnaviridae*, *Smacoviridae*, and *Sphaerolipoviridae* showed positive correlations with poultry servings, meat servings, and refined grain servings, respectively (**Fig. 3A**). Furthermore, we noted that certain demographic factors, including the age at the diagnosis of MS, demonstrated a significant positive correlation with the relative abundance of *Myoviridae*. In contrast, this demographic factor showed negative correlations with the relative abundance of *Bicaudaviridae* and *Alphaflexiviridae* (**Fig. 3A**).

**Fig. 3.**
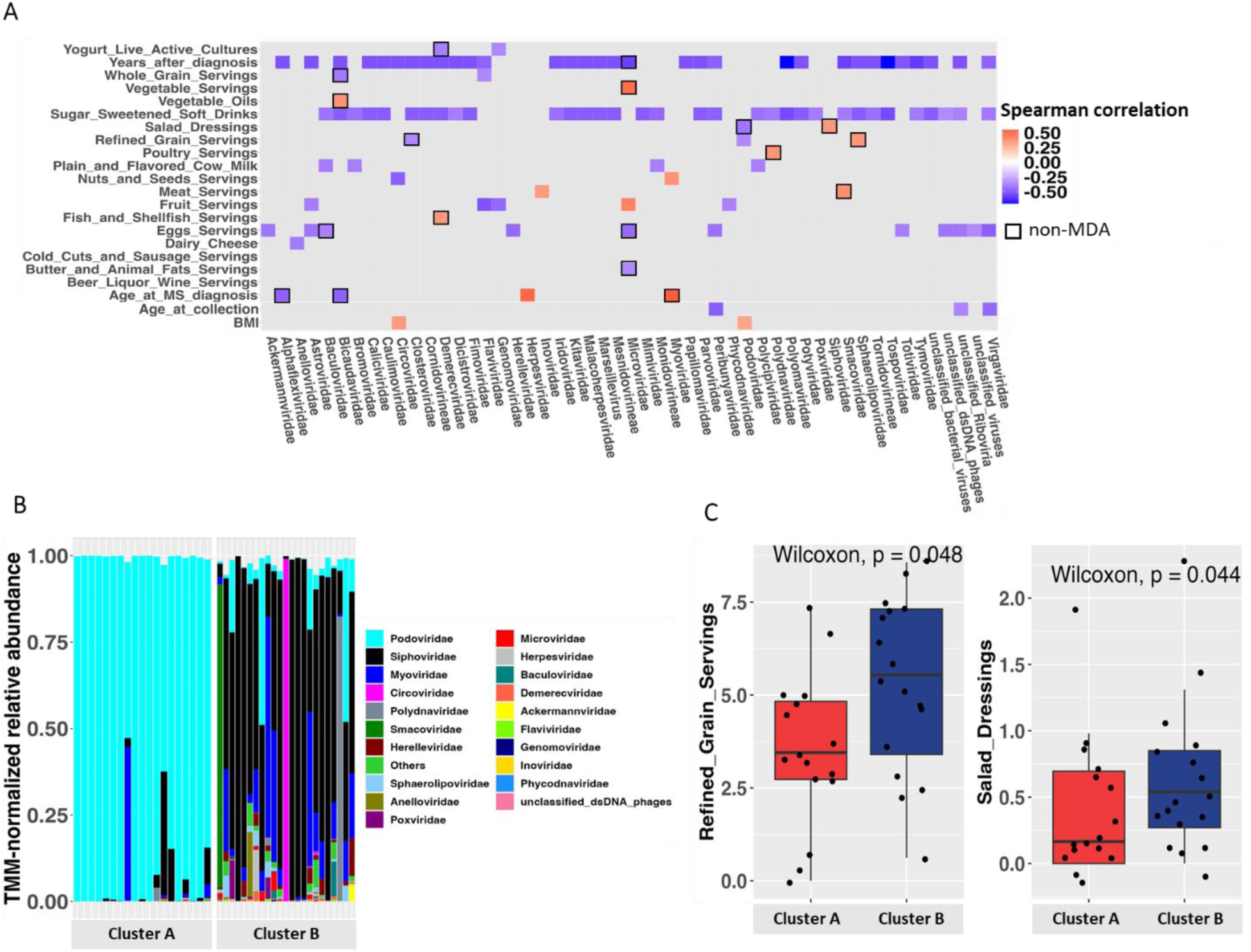
presents an analysis of the associations between diet, demographic characteristics, and clinical factors with the abundance of viruses. **A**: The heatmap illustrates Spearman’s correlation of the relative abundance of different gut viruses and dietary habits within both RRMS and HC groups. The size and color intensity of the squares in the heatmap indicate the strength of correlation, with blue squares representing negative correlations and orange squares representing positive correlations. Correlations that are statistically significant are highlighted with a black square border; **B**: A stacked bar plot displays the top 20 most abundant gut viruses between the *Podoviridae*-dominated cluster A and the *Siphoviridae*-dominated cluster B; **C**: A box plot visualizes a significant dietary factor comparison between the *Podoviridae*-dominated cluster A and the *Siphoviridae*-dominated cluster B. The box plot shows the data distribution, with the median indicated by a horizontal line within the box. The “box” encompasses the middle 50% of the data, while the upper and lower whiskers represent the maximum and minimum values, respectively. Statistical comparison of dietary factors was conducted using a Wilcoxon test, with corresponding *p*-values provided.

In our study, we observed a distinct virus relative abundance pattern among non-MDA samples with one subset being predominantly dominated by the viral family *Podoviridae* and another subset showing a lesser but noticeable dominance by *Siphoviridae*. We performed a cluster analysis for the gut virome at the family level and identified two virome clusters, labeled as Cluster A and Cluster B, primarily based on the prevalence of these two viral families (**Fig. 3B**). Within Cluster A, we found that 17 out of 24 subjects had relative abundance of *Podoviridae* exceeding 80%, with 15 of those subjects having relative abundance greater than 90%. Conversely, in Cluster B, 14 out of 18 subjects displayed *Siphoviridae* relative abundance exceeding 50%, with 5 of those subjects having relative abundance greater than 90%. Notably, Cluster B, which was dominated by *Siphoviridae*, exhibited a more diverse composition of viral families. When comparing the dietary metadata between these clusters, we found that subjects in Cluster B, which was dominated by *Siphoviridae*, had significantly higher servings of refined grains (*p*=0.048), and salad dressings (*p*=0.044), in contrast to those in Cluster A, which was dominated by *Podoviridae* (**Fig. 3C**). Interestingly, constipation was reported by nineteen subjects in our cohort, and they all belonged to Cluster A, with no instances reported in Cluster B, showing a statistically significant difference (*p*=0.027). Furthermore, subjects in Cluster A exhibited significantly higher BMIs (*p*=3.1E-06).

### Correlation between MS-associated bacteria and viruses

To understand association between the gut virus and bacteria, we constructed a correlation network (**Fig. 4**). This network draws connections between the presence of various bacterial genera and viral families in a combination of RRMS and HC samples. In this network, there were a total of 49 nodes, which represented 15 viruses and 34 bacteria. These nodes were connected by 57 edges, representing 32 positive correlation and 25 negative correlations. For instance, *Siphoviridae* exhibited interactions with 5 bacterial genera, which was the highest number of correlation (all positive correlation) observed for any single phage in the network. In contrast, *Microviridae* displayed highest negative correlation with 5 bacteria genera. This suggests that targeted removal or addition of these two phages could potentially induce more significant alterations in the bacterial community compared to a virus like *Poxviridae*, which only interacted with one bacterial genus. Furthermore, network analysis revealed that *L. bacterium* and *B. longum*, two bacterial genera, were involved in the highest number of correlations with four and three viruses, respectively. These bacteria were also predicted to interact with *Siphoviridae* and *Microviridae*.

**Fig. 4.**
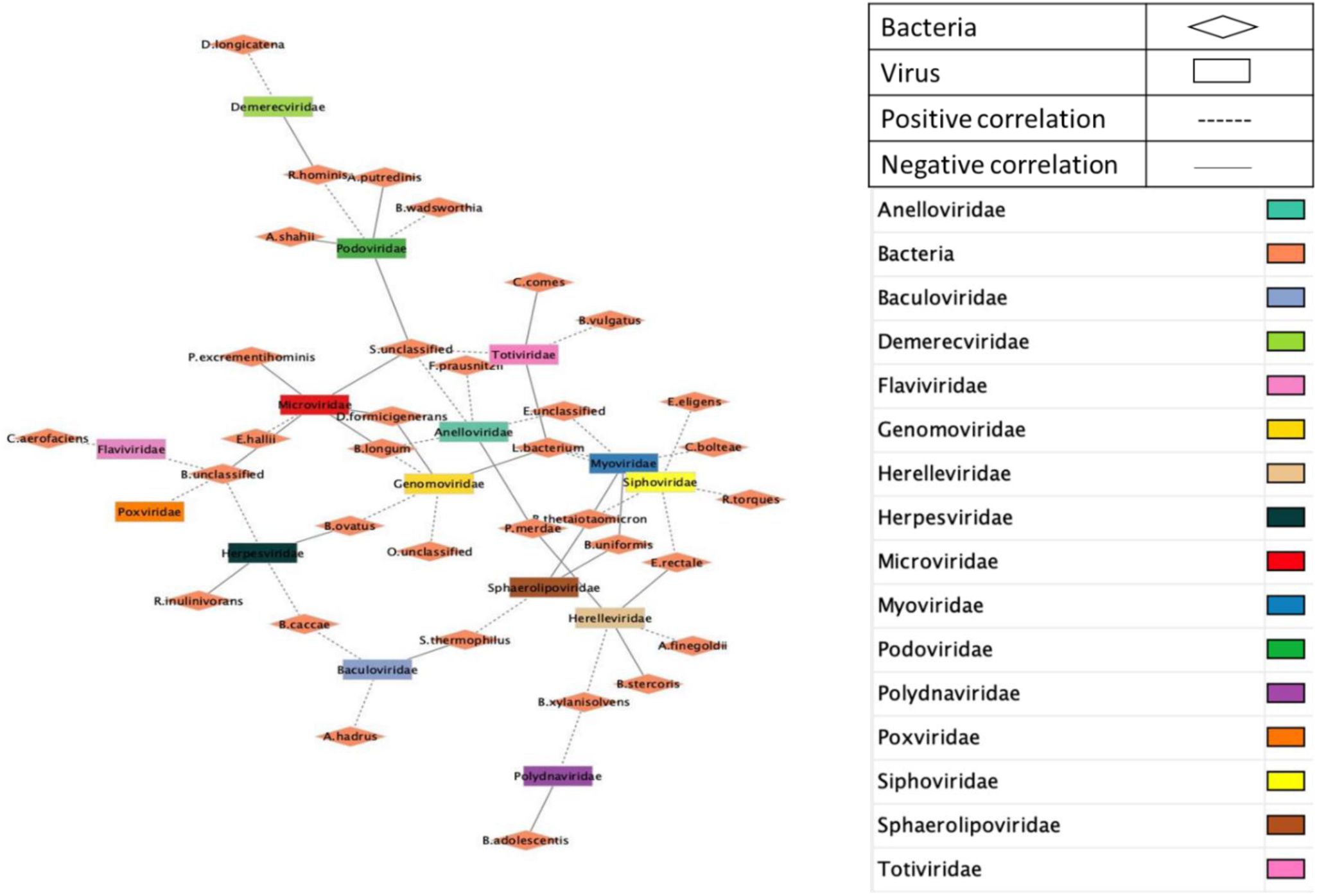
displays a network plot illustrating the relationships among bacterial genera and viral family within the gut microbiome of individuals with RRMS. Each node in the plot represents either a virus family (rectangle) or a bacterial genus (diamond). Edges connecting nodes such as dashed lines indicate significant positive correlation, while solid lines represent significant negative correlation.

### crAss-like phage in MS

Our analysis revealed the consistent presence of crAss-like phage in both methods. In non-MDA samples, crAss-like phage dominated approximately 50%, while it comprised a slightly lower proportion (~40%) in MDA samples (**Fig. 5A**). Notably, a significant (*p*=0.044) increase in the relative abundance of crAss-like phage was observed in RRMS in non-MDA method (**Fig. 5B**). Exploring virus–bacterium associations, we found a positive correlation between the relative abundance of *Bacteroides* and crAss-like phage. However, the distribution of their relative abundance was unexpected (R=0.35 and *p*=0.025) (**Fig. 5C**). In certain samples with reduced *Bacteroides*, there was a notable abundance of crAss-like phages, whereas samples with highly abundant *Bacteroides* exhibited no detectable crAss-like phage. This suggests that host bacteria in the latter case might have shed their receptors essential for the docking and entry of crAss-like phages. Notably, *B. fragilis*, acting as a host for crAss-like phage, showed a positive correlation (R = 0.41 and p < 0.01) with crAss-like phage (**Fig. 5D**). On the other hand, other *Bacteroidetes*, such as *B. intestinalis*, also serving as a host for crAss-like phages, displayed a non-significant correlation (R = −0.014 and p = 0.93). Considering the potential family-scale taxonomic diversity of crAss-like phages, it is plausible that they may infect a broad range of hosts within *Bacteroidetes* species, resulting in weak relative abundance correlations with specific host taxa.

**Fig. 5.**
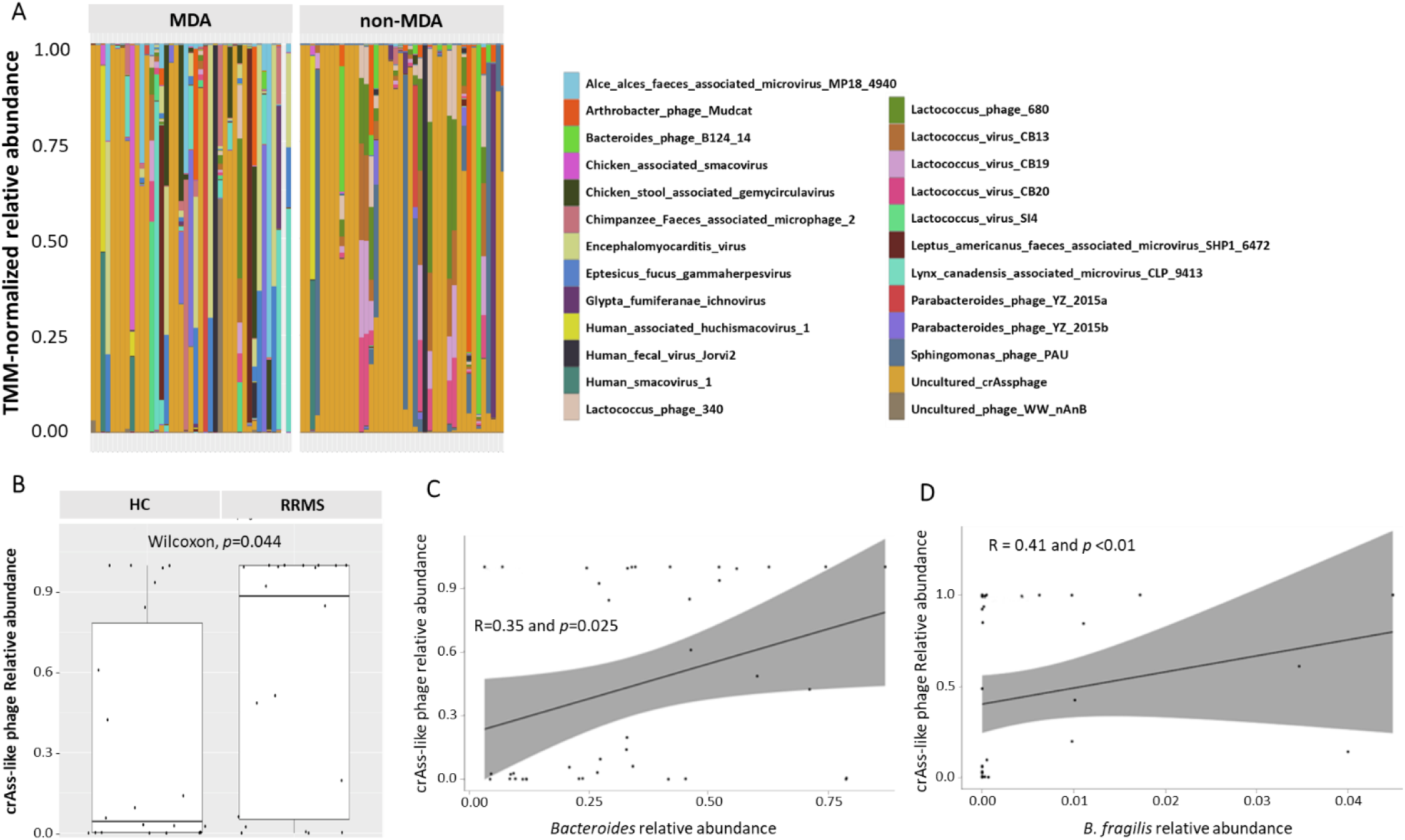
abundance and correlation of crAss-like phage with host bacteria **A**: A stacked bar plot displays the top 20 most abundant gut viruses at organism level in both MDA and non-MDA methods; **B**. crAss-like phage relative abundance in RRMS and HC. Statistical comparison was conducted using a Wilcoxon test, with corresponding *p*-values provided; **C** and **D** Correlation between crAss-like phage abundance and host bacteria. Correlations that are statistically significant are provided in panel.

## Discussion

The increasing adoption of VLPs enrichment technology, coupled with high-throughput sequencing methods, has brought significant focus to virome research, especially in the context of DNA virome. In our study, we extracted VLP genomic DNA and performed deep metagenomic sequencing with and without MDA amplification to profile the gut virome in individuals with RRMS and HC. In this study, we revealed the alteration in gut virome in RRMS patients compared with HC. MDA is known for its significant impact on reducing genetic diversity and reproducibility, often over-representing single-stranded DNA viruses (39). Despite observing no significant difference in overall virome evenness, our study revealed that MDA samples exhibited a lower viral hit rate and viral richness compared to non-MDA samples in both RRMS and HC subjects. MDA is also known for its lower error rate due to exonuclease and proofreading activities (40). It can generate long amplification products (50-100 kb), making it suitable for long-read sequencing and whole-genome sequencing (41). However, both MDA and non-MDA methods discovered large number of non-overlapping viruses, thus allowing us to have a comprehensive view of the gut microbiome in these participants.

Our study further delves into the virome of individuals with RRMS. The findings revealed a significant decrease in gut virome diversity in RRMS patients compared to HC, which aligns with previous studies in IBD and autoimmune diseases (17, 42). Considering that bacteriophages constitute the majority of the gut virome, the reduced richness and diversity of the gut virome in RRMS patients may be indicative of a scarcity of bacterial hosts (43). The observed dysbiosis in the gut virome further showed equal importance to bacterial diversity in response to changes in the gut microbiome associated with RRMS. Further, MDA and non-MDA revealed distinct differences in the distribution and composition of viral assemblages. The non-MDA method resulted in a more even distribution of double stranded DNA viral assemblages and a higher number of rare groups compared to the MDA methods. Additionally, non-MDA predominantly led to a viral composition dominated by bacteriophage Caudovirales order, including *Siphoviridae*, *Myoviridae*, and *Podoviridae* (26, 44). In contrast, MDA samples exhibited a greater proportion of circular, single-stranded DNA viruses from the *Microviridae* family, as well as enhanced proportions of eukaryotic viral families *Genomoviridae*, *Circoviridae*, and *Anelloviridae*. These findings highlight the influence of the viral enrichment method on the diversity and distribution of the detected viral elements. The differences observed between the MDA and non-MDA results could be attributed to variations in denaturation procedures, MDA kit vendors, or template amounts during random priming (45). However, it is important to note that random amplification bias has minimal impact on inter-subject beta diversity due to the uniqueness of human gut virome (45).

Following filtration for contamination and low-relative abundance sequences, we observed a significant decrease in *Siphoviridae* and *Myoviridae* within the metagenome of RRMS patients, while *Podoviridae* and *Microviridae* were more prevalent in RRMS compared to HC. *Siphoviridae* were skewed towards HC. A recent report from Japanese patients indicates a nominal increase in abundance of *Podoviridae* and *Microviridae*, coupled with a decrease in the abundance of *Siphoviridae* and *Myoviridae* viral families in MS patients. (26). This suggests that variations in gut virome may be attributed to differences in viral enrichment approaches or population-specific factors, emphasizing the need for further investigation in future experiments. Our findings suggest a potential association between *Podoviridae* and *Microviridae* relative abundance and MS pathogenesis. These gut virome families were more abundant in RRMS, though their biological relevance in health and disease requires further exploration. Bacteriophages can indirectly influence the host’s immune system through their interactions with bacteria. Specifically, *Siphoviridae*, with a significantly increased relative abundance in HC, showed the highest number of positive correlations. Conversely, *Microviridae*, with a significantly increased relative abundance in individuals with RRMS, exhibited the highest number of negative correlations. These findings suggest that the gut virome may exert its impact on MS by regulating bacterial populations and underscoring the need for further research to unveil the intricate interplay between the bacteriome, bacteriophages, and MS.

Interestingly, the sole dietary factor that significantly differed between RRMS patients and HC was meat consumption (46), which showed a positive correlation with *Smacoviridae*. Our dietary pattern analysis of RRMS and HC indicated that increased meat consumption might be associated with a higher risk of developing multiple sclerosis. We speculated that alterations in the microbiome, particularly shifts in the virome linked to meat consumption, may initiate a cascade of events contributing to or exacerbating the disease. This dietary pattern has the potential to unveil masked connections that could enhance our knowledge of the disease’s etiology. Despite the crAss-like phage being commonly present in a healthy gut virome, many of its proteins do not share similarities with previously identified viral proteins (47). The connection between crAss-like phages and disease states, especially autoimmune diseases, has been an area of interest (26). In our investigation, we observed a significant increase relative abundance of crAss-like phages in RRMS patients. Tomofuji et al. (26) reported increased crAssphage abundance in MS, although the difference was not statistically significant. Conversely, crAssphage abundance was found to be reduced in other autoimmune disorders such as systemic lupus erythematosus and rheumatoid arthritis (26). Further, crAss-like phages were also skewed in viral families dominated by *Podoviridae*, a skewness that was significant in RRMS. We also identified a positive correlation between host *Bacteroides* and crAss-like phages. However, the distribution of their relative abundance was not normally shaped, suggesting a probable family-scale taxonomic diversity among crAss-like phages. This diversity may allow them to infect a wide range of hosts across *Bacteroidetes* species, resulting in poor relative abundance correlations between crAss-like phages and specific host taxa. These findings collectively point to disease-specific gut virome and bacterial microbiome in MS. The findings regarding crAss-like phage further emphasize the complexity of the gut virome and its potential associations with MS. Future research is warranted to unravel the intricate relationships between the gut bacteriome, virome, and MS pathogenesis, which could open up new avenues for therapeutic interventions and personalized medicine approaches.

## Limitation

The study has several limitations that need to be acknowledged. One limitation is the sample size of MS patients in our study compared to the general population. Consequently, conducting larger-scale analyses with increased statistical power is essential to thoroughly investigate the association between the gut virome and MS. Additionally, it is worth noting that the majority of RRMS patients in our study were in remission, with only a few receiving treatments. Another limitation pertains to the bias in enrichment associated with the MDA and non-MDA method. Careful consideration is required when selecting amplification or not in sequencing library preparation. The sequencing process resulted in numerous unclassified viruses, further underscoring the study’s limitations.

## Conclusion

Our comprehensive case-control study has illuminated a previously undisclosed facet of the relationship between the gut virome and MS. Analysis of the taxonomic microstructure in RRMS and HC has unveiled distinctive variations in the specific bacteria and viruses identified. These findings hint at a novel perspective wherein RRMS may be associated with an increase in bacteriophage diversity, primarily linked to their bacterial host cells. It is noteworthy that we observed both positive and negative correlation between specific virus and bacterial taxa, suggesting that these alterations could potentially play a role in disease pathogenesis. To gain a more comprehensive understanding of the dynamics of the gut virome in MS, future research should encompass a larger sample size and consider cross-sectional and longitudinal time points. Such an approach will facilitate the exploration of the intricate processes underlying the observed changes, including their associations with diet, disease development, and other potential factors.

## Declarations

### Ethics approval and consent to participate

Ethical approval was obtained prior to the study.

### Availability of data and material

Upon a reasonable request, all datasets will be available from the corresponding author.

## Acknowledgments

We thank The Jackson Laboratory for Genomic Medicine, CT, U.S.A. for sequencing.

